# Rigorous Quantitative Analysis of Nonlinear Uncertain Biomolecular Systems using Validated Methods

**DOI:** 10.64898/2025.12.13.693835

**Authors:** Rudra Prakash, S. Janardhanan, Shaunak Sen

**Author notes:** Corresponding author: Rudra Prakash.

## Abstract

The paper addresses the critical challenge of accurately characterising steady states in biomolecular systems, which are often complex, nonlinear, multistable and subject to significant uncertainties. Traditional numerical methods often fail to provide complete or guaranteed solutions under these conditions. To overcome these limitations, the research proposes and evaluates the application of interval analysis methodologies. We provided algorithms for interval Newton and interval Krawczyk methods for rigorously bounding all possible steady states (both stable and unstable) in multistable, multidimensional nonlinear systems. This study involves a comparative analysis of these two methods in conjunction with interval bisection and interval constraint propagation. We addressed numerical examples for an array of biologically plausible models, involving both feedback and feedforward gene networks. The work recommends the choice of the most suitable method for various types of biomolecular systems, ultimately offering a robust computational framework to understand cellular functions and design synthetic biological circuits.

## 1. INTRODUCTION

Biomolecular systems are characterised by nonlinear dynamics, multistability, multi-dimensionality, and considerable uncertainty, and each poses significant challenges in the fields of drug design and synthetic biology prediction and control (Del Vecchio and Murray, 2015; Alon, 2006). For example, nonlinear signalling cascades such as the MAPK/ERK pathway show ultrasensitive dynamics, which complicates drug efficacy prediction (Filippi et al., 2016). Multistable apoptosis networks, regulated by Bcl-2 interactions, fluctuate between survival and death, challenging therapeutic control (Bagci et al., 2006). Synthetic circuits, such as toggle switches (Gardner et al., 2000), feedback and feedforward loops (Alon, 2006; Ma et al., 2009), rely on steady-state properties to enable functions such as switching, adaptation or signal processing. In biomolecular systems, such multistability is crucial for cellular processes, including differentiation, apoptosis, and phenotypic switching (Angeli et al., 2004; Ferrell, 2002). Nonlinear dynamical systems often exhibit multiple steady states, each linked to distinct long-term behaviours. In practical applications, safe and reliable steering of cellular processes and the development of molecular therapies depend not only on the search for possible stable solutions but also on rigorously bounding these solutions under real-world uncertainty.

Accurately identifying and characterising steady states is essential not only for understanding the operation of natural regulatory systems but also for the construction of synthetic networks.

Numerical simulations, parameter sweeps, and bifurcation analyses are predominantly employed in contemporary research within systems and synthetic biology to identify multiple steady states (Angeli et al., 2004; Hasenauer et al., 2009; Otero-Muras et al., 2012; Kuntz et al., 2019; Sreedharan et al., 2023; Reyes et al., 2022; Avcu and Güzeliş, 2019; Dey and Barik, 2021). Although these techniques are practically effective, they do not ensure completeness or guaranteed validation. Traditional numerical solvers, for instance, might fail to detect equilibria, produce erroneous solutions, or become unreliable under parameter uncertainty or in the presence of an unstable equilibrium point. This limitation holds significant implications in biology, where minor variations in kinetic rates may result in divergent cellular outcomes, and in engineering, where unforeseen equilibria could compromise safety and stability. Consequently, there is a burgeoning demand for computational methodologies that provide mathematically rigorous and guaranteed bounds on the steady states of nonlinear multidimensional systems.

Validated numerical methodologies assure these certainties. Interval analysis provides a means to represent uncertain quantities as intervals rather than single values (Moore et al., 2009). By propagating these intervals through interval arithmetic operations, we can obtain enclosures that rigorously contain the true solution set. Several interval analysis-based algorithms have been proposed for calculating steady-state bounds (Neumaier, 1990; Jaulin et al., 2001; Rump, 2010). These methods have been effectively employed within cyber-physical systems, robotics, and chemical process systems to determine steadystate bounds in the presence of parameter uncertainty (Hasenauer et al., 2009; Jaulin et al., 2001; Tucker, 2011). However, interval analysis is not widely used for the analysis of biomolecular networks. Addressing this gap is crucial, as rigorous steady-state analysis would prevent biologically erroneous conclusions and offer reproducible assurances independent of solver or initial condition choices. Validated numerical methods can calculate reliable bounds on steady-state concentrations, offering insight into the robustness of the system and its sensitivity to parameter changes.

Previously, we effectively employed the Interval Newton method, which is an interval-based adaptation of the traditional Newton method, to achieve steady-state enclosures in models of multistable scalar and monostable two-dimensional biomolecular systems (Chorasiya et al., 2023). This study focuses on solving steady states for complex multidimensional and multistable biomolecular systems in the presence of parametric uncertainties.

The main contributions of this paper are summarised as follows. First, we present algorithms pertinent to the Interval Newton Method and the Interval Krawczyk Method for the computation of all potential roots within multidimensional, multistable non-linear systems with uncertain parameters. Second, we perform a comparative analysis of four methods grounded in interval analysis: (*i*) interval bisection, (*ii*) interval constraint propagation, (*iii*) interval Newton, and (*iv*) interval Krawczyk. Third, our study offers recommendations regarding the suitability of the presented methods for various types of problems. The rationale behind this is that no single interval method represents an optimal solution for all types of problem. Interval methods are compared based on their overall rigour, efficiency, robustness to different problems. And fourth, the methodology is demonstrated through representative synthetic and natural motifs, including negative feedback loops, feedforward loops (Ma et al., 2009; Del Vecchio and Murray, 2015; Kim et al., 2014), and multistable gene networks (Alon, 2006; Venturelli et al., 2012).

The organisation of this paper is as follows: Section II offers a brief overview of interval analysis. In Section III, we describe a systematic stepwise approach for applying the interval methods. Section IV illustrates examples of biomolecular systems alongside the outcomes of our comparative analysis, emphasising the advantages and limitations of each algorithm. Finally, Section V concludes with a summary of our findings and suggestions for future research directions.

## 2. METHODOLOGY

### 2.1 Preliminaries

#### Interval Analysis

Interval analysis constitutes a rigorous method for representing quantities using intervals instead of single point values. This method’s unique characteristic is its ability to encode both the actual value and its inherent uncertainty concurrently. Fundamentally, interval analysis is concerned with intervals of real numbers, each defined by a distinct lower and upper bound. Consider two intervals, 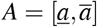 and 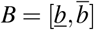, where 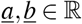 and 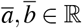 represent the lower and upper bounds, respectively, with *A* and *B* belonging to the set of all intervals, denoted as 𝕀 ℝ. Arithmetic operations, including addition, subtraction, multiplication, and division, are generalized to operate on these intervals, producing result intervals that encompass all conceivable outcomes of the associated operations on real numbers contained within the input intervals. This essential attribute ensures that interval computations inherently encompass the true solution, regardless of existing uncertainties. The operations of interval arithmetic can be articulated through the following table.

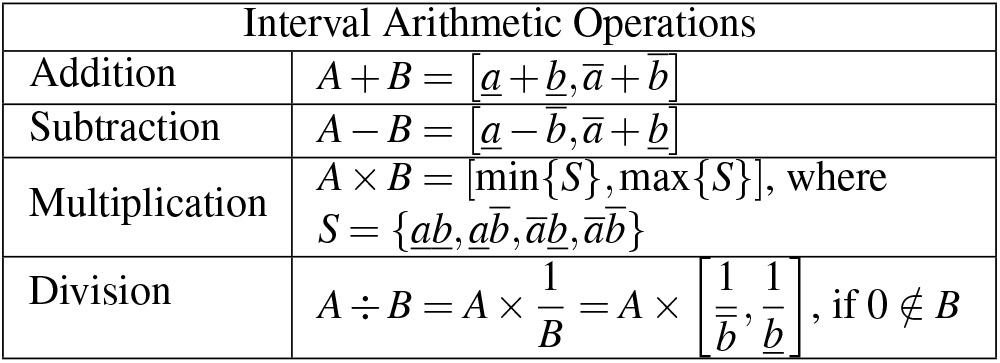

Extended interval arithmetic is applied when dividing by an interval that includes zero. The *n*-dimensional intervals can be expressed as the Cartesian product of all individual *n* intervals, commonly referred to as an “Interval Box”. Beyond fundamental arithmetic, interval analysis encompasses functions, allowing the assessment of functional ranges over intervals of input values. The subsequent theorems underscore two pivotal properties in the interval-based analysis: inclusion isotonicity and range enclosure.

##### Theorem 1.

(Moore et al., 2009) Given a real-valued, rational function *f* and a natural interval extension *F* such that *F*(*X*) is well defined for some *X* ∈ 𝕀ℝ, we have

1. *Y* ⊆ *Z* ⊆ *X* ⇒ *F*(*Y*) ⊆ *F*(*Z*), (*inclusion isotonicity*)
2. *R*(*f* ; *X*) ⊆ *F*(*X*), (*Range Enclosure*).

Despite its advantages, interval analysis encounters certain difficulties, particularly concerning the overestimation of bounds. This issue largely stems from the interval dependency problem, which occurs when a variable appears multiple times within an expression. Utilizing the theorem provided here, one can ensure that the range is enclosed with the desired precision.

##### Theorem 2.

(Moore et al., 2009) Consider *f* : *I* → ℝ with *f* Lipschitz and let *F* be an inclusion isotonic interval extension of *f* such that *F*(*X*) is well-defined for some *X I*. Then there exists a positive real number *k*, depending on *F* and *X*, such that, if 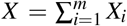, then 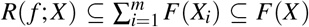 and rad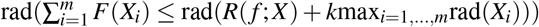

### 2.2 Interval Methods

The properties of interval analysis previously discussed can be employed to identify the solution region, if a solution exists, within a specified search space.

#### Interval Bisection Method

The interval bisection method is an interval version of the conventional Bisection technique to find the root(s) of a system. This method operates by recursively dividing intervals into sub-intervals until a specified condition is met or a desired level of precision is achieved. This iterative process involves systematically bisecting intervals and evaluating the function within each sub-interval. As compared to the standard bisection method, the interval version is capable of finding all roots of *f*. The stepwise procedure to calculate interval root(s) is given below.

***Step 1:*** Assume a sufficiently large initial enclosure of the steady state(s), *X*_0_.

***Step 2:*** Check if 0 ∈ *F*(*X*_0_,*U*). If *true*, then **Bisect** *X*_0_ into *right* (*X*_*r*_) and *left* (*X*_*l*_) halves. If *false* then discard *X*_0_.

***Step 3:*** Follow **Step 2** recursively until width of the bisected halves is less than given *tolerance* (*δ*) value.

The initial domain are either proved not to contain any roots, whereby they are discarded from the search or are bisected and kept for further study. When all remaining subintervals are smaller than some predefined tolerance, the search is terminated. We are then left with a collection of intervals whose union covers all possible roots of *f* within *X*_0_.

The Interval Bisection method, while straightforward to implement for locating multiple roots of the equation *f* (*x, u*) = 0, presents two notable drawbacks. First, it necessitates subdividing the entire search domain until the tolerance condition is met, rendering it computationally expensive for higher-dimensional systems. Second, in systems exhibiting multiple steady states, a sufficiently small tolerance value must be chosen to ensure that no two roots reside within the same interval box, further contributing to its computational cost.

Consequently, an alternative approach to eliminate recursion is the Subdivision + Filter method. This method subdivides the search space and discards subboxes that do not meet the existence criteria, thereby identifying those that do not contain a solution (Prakash et al., 2024).

The Interval Constraint Propagation method offers an advantage in handling multiple steady states and higher-dimensional systems efficiently.

#### Interval Constraint Propagation

Interval Constraint Propagation is a computational technique used to solve constraint satisfaction problems involving continuous variables. Its aim is to reduce the search space for potential solutions by progressively narrowing the intervals of variables based on the specified constraints. This reduction is achieved through the concept of set inversion, which can be illustrated by the following theorem.

##### Theorem 3.

(Jaulin, 2000) Assume that it is possible to isolate *u*_*i*_ in the expression *f* (*x, u*) = 0, i.e., there exists a function *g*_*i*_ that satisfies *f* (*x, u*) = 0 ⇔ *u*_*i*_ = *g*_*i*_(*x*, ^*i*^*u*), where ^*i*^*u* = (*u*_1_, …, *u*_*i*_ −_1_, *u*_*i*+1_, …, *u*_*n*_)^*T*^. The function *g*_*i*_ is called the solution function associated with *u*_*i*_. Denote by *π*_*i*_ the projec-tion operator on the *i*^*th*^ axis and by *G*_*i*_ and inclusion function for *g*_*i*_. We have *π*_*i*_(*s*) ⊂ *G*_*i*_(*X*, ^*i*^*U*) ∩ *U*_*i*_, where ^*i*^*U* = (*U*_1_, …,*U*_*i* −1_,*U*_*i*+1_, …,*U*_*n*_)^*T*^. Moreover, if *g*_*i*_ is continuous and if *g*_*i*_ is minimal, then the inclusion becomes an equality.

The aforementioned theorem can be applied to equations with complex constraints by adhering to the following procedure.

To solve the problem, begin by decomposing the main constraint into individual primitive constraints and construct the forward contractor. Subsequently, construct the backward contractor by deriving it from the forward contractor but in reverse order, starting from the end and moving to the beginning. At each stage within a primitive constraint, ensure that each variable on the right-hand side is isolated appropriately. Following this, proceed to intervalise both the forward and backward contractors. These combined intervalised algorithms form what is collectively termed a ‘contractor’. Lastly, integrate these intervalised algorithms to complete the solution process (Jaulin et al., 2001; Prakash et al., 2024).

Interval constraint propagation is capable of handling systems with multiple constraints simultaneously, making it suitable for addressing problems with complex nonlinearity.

We have proposed algorithms related to the following two methods, which use information regarding a function’s derivative. As a result, this improves convergence efficiency.

#### Interval Newton Method

The Interval Newton Method is a numerical technique employed to determine the roots of functions with real or interval values within a specified interval. It extends the classical Newton-Raphson method to accommodate interval arithmetic, rendering it especially advantageous to address parametric uncertainties or locate multiple roots within an interval box (Chorasiya et al., 2023). Algorithm 1 is designed to calculate multiple bounded solutions to a nonlinear system of equations.

##### Algorithm 1

Interval Newton Method

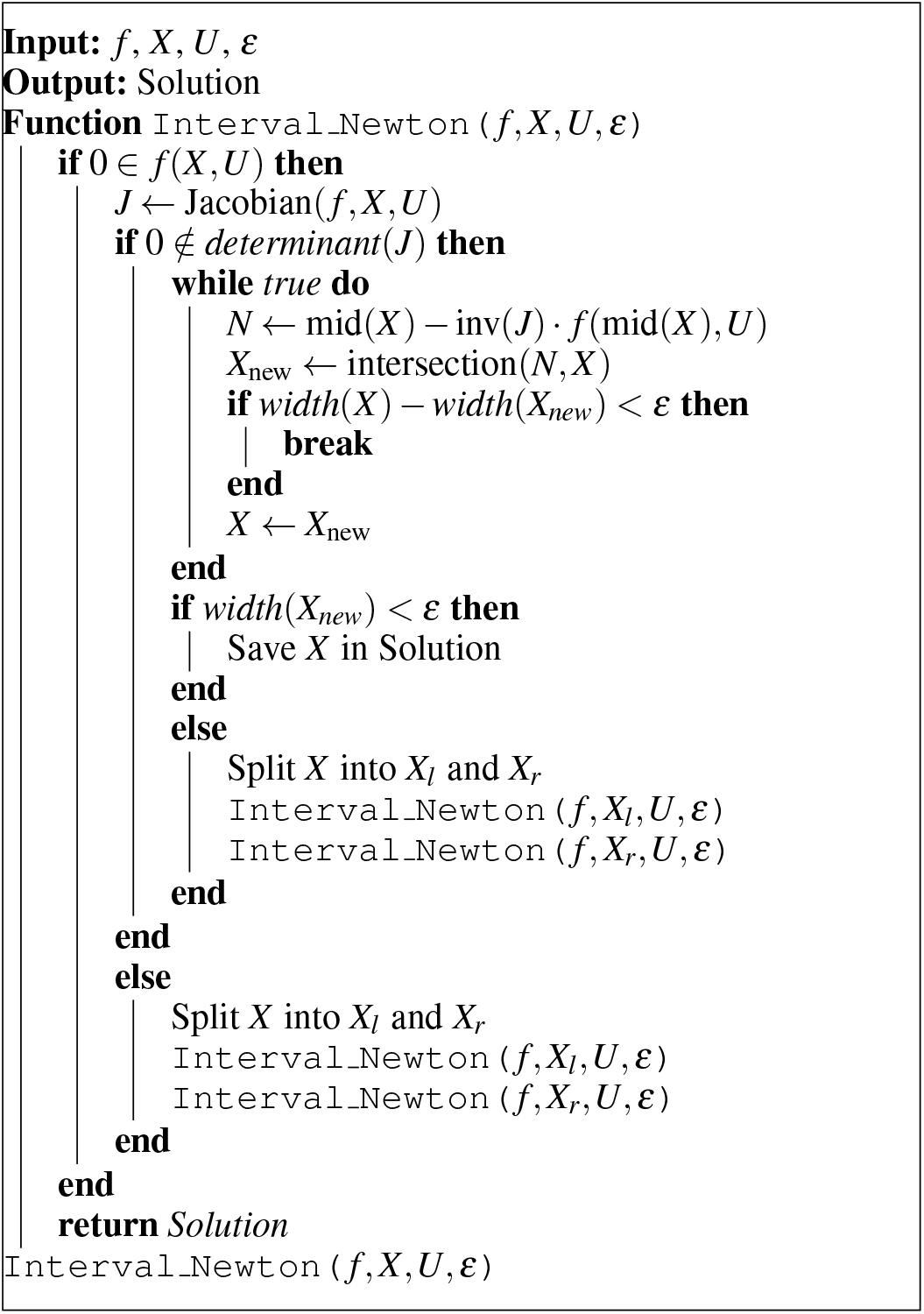

In systems exhibiting multiple steady states, where 0 ∈ *F*_*x*_, extended interval arithmetic is employed. The Interval Newton Method is advantageous due to its rapid convergence speed, attributable to its quadratic convergence rate. However, in a system of equations, the method necessitates division by an interval Jacobian matrix. This presents a challenge in ill-conditioned systems where the Jacobian matrix is singular, thus hindering convergence. The Interval Krawczyk Method offers an alternative approach to effectively address this issue.

#### Interval Krawczyk Method

The Krawczyk method is another version of the interval Newton method, distinctively characterised by its ability to operate without the need for the inversion of an interval matrix (Moore, 1977). Algorithm 2 illustrates the implementation of the Krawczyk method for bounding solutions in multistable systems.

##### Algorithm 2

Interval Krawczyk Method

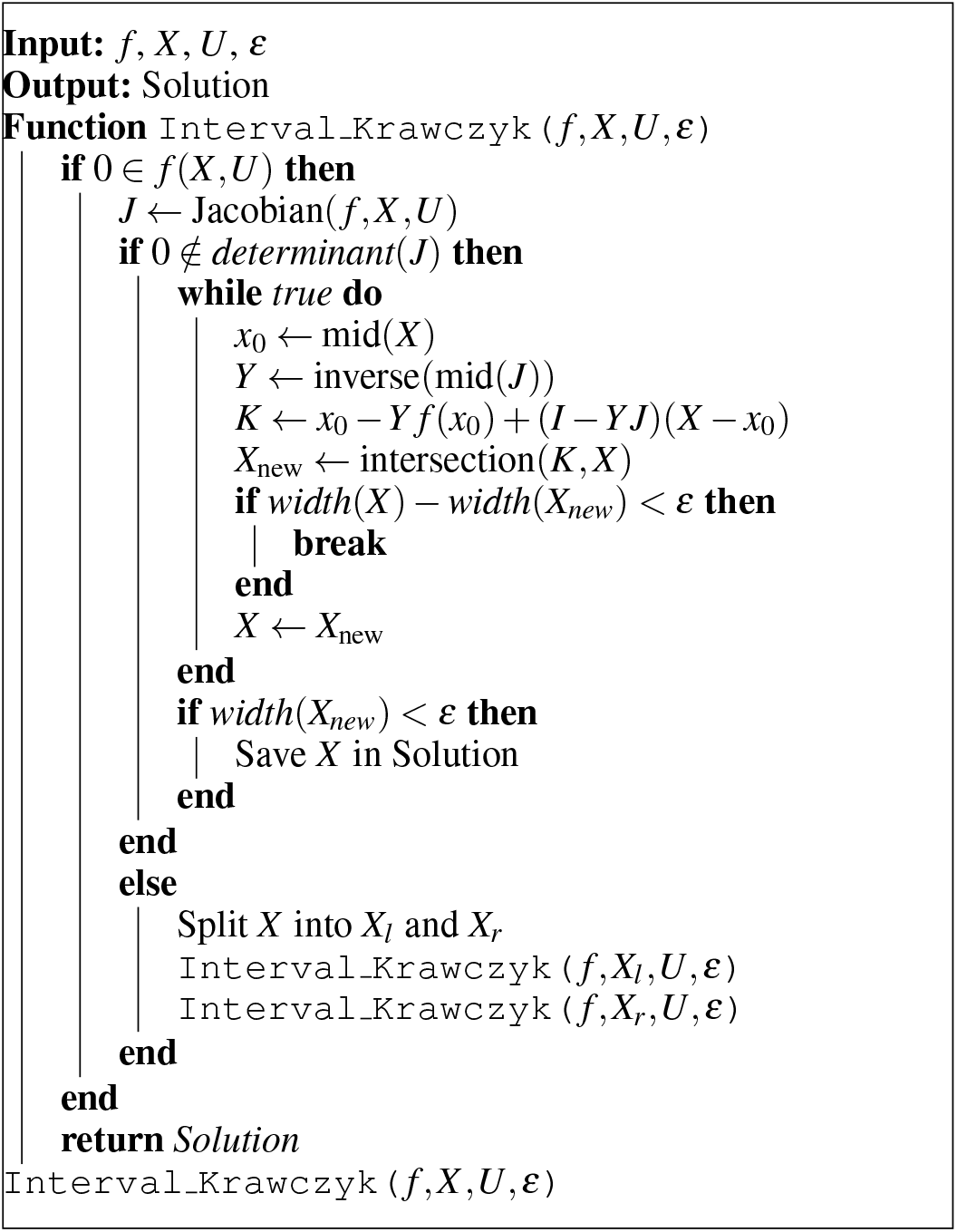

The Interval Krawczyk Method, while exhibiting relatively slower convergence compared to the Interval Newton Method, demonstrates superior robustness when applied to ill-conditioned multidimensional systems.

For functions involving interval parameters, such as the Newton operator *N*(*X,U*) and the Krawczyk operator *K*(*X,U*), the resulting interval solutions tend to overestimate the solution bounds. This overestimation can be mitigated through the subdivision of interval parameters (Chorasiya et al., 2023).

##### Corollary 1.

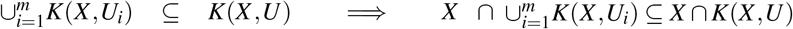

## 3. RESULTS & DISCUSSION

To study and compare the underlying interval methods, we study some well-known biomolecular systems comprising various levels of complexity, single or multiple steady states, and scalar or multidimensional biomolecular systems. The computations were implemented in Julia 1.11.6 with the packages IntervalArithmetic v0.20.3, ReversePropagation v0.2.3, and Symbolics v5.28.0. The code has been available on GitHub (Exa, 2025).

### 3.1 Negative Feedback Loop

Negative feedback loops constitute one of the principal network motifs in adaptive biomolecular systems, serving critical regulatory functions. Within this nonlinear model, the protein synthesis process is shown to exhibit self-inhibition. The mathematical representation of this dynamical model is described below:

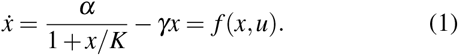

Here, *x* denotes the concentration of the protein, while *α, γ*, and *K* signify the maximal protein production rate, the rate constant for degradation, and the DNA binding constant of the protein to its own promoter, respectively. These parameters are conveniently expressed in the parameter vector *u* = [*α, γ, K*]^*T*^. For analysis purposes, the parameter values *α* = [200, 300], *γ* = [2, 3], and *K* = [200*/*3, 150] are considered, with the initial enclosure of the solution represented by *X*_0_ = [0, 500]. The bounded steady states of the negative feedback loop, using the interval bisection method, are depicted in Figure 1.

**Fig. 1.**
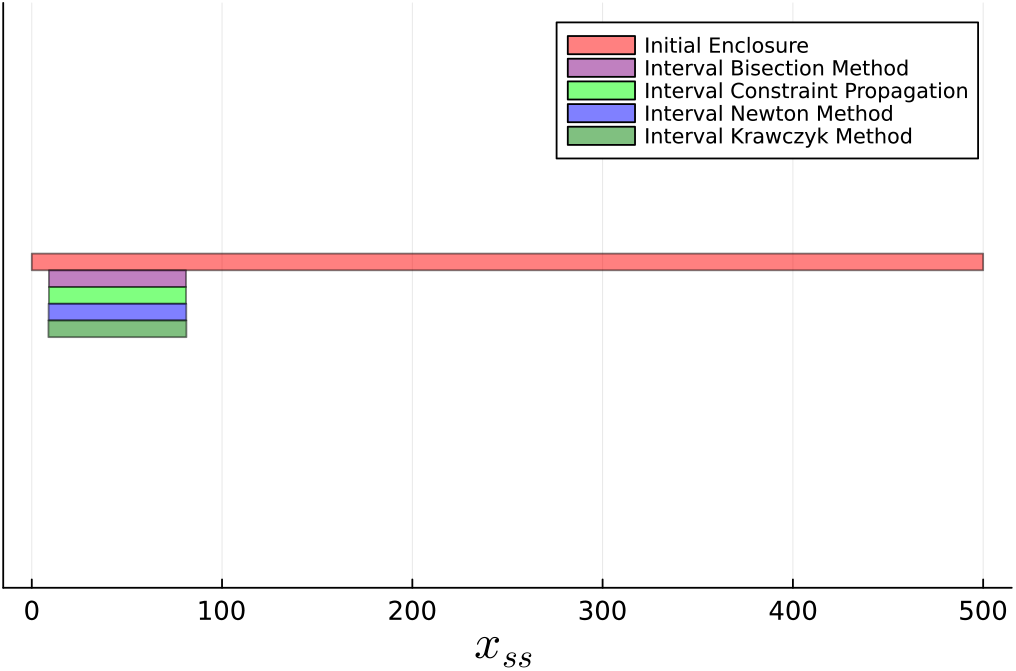
Negative Feedback Loop: Rigorous bounds on steady state for given parameter interval using interval methods. The parameter ranges under consideration are *α* = [200, 300], *γ* = [2, 3], and *K* = [20*/*3, 15]. The initial condition for the solution is denoted by the interval *X*_0_ = [0, 500].

### 3.2 Positive Feedback Loop

The positive feedback loop constitutes a crucial component in biomolecular systems by amplifying protein production through self-activation. This model is pivotal for interval-based analysis due to its characteristic multi-steady-state nature, offering insights into complex biological processes.

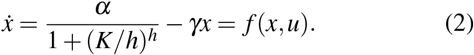

The Hill coefficient *h* is a parameter that describes the cooperativity of ligand binding in biochemical systems. Its value affects the sensitivity of the response in systems governed by the Hill equation.

This serves as a robust example to illustrate the limitations of traditional root-finding techniques, like the Newton-Raphson method. The Newton-Raphson method is known for its local convergence, yet it has several limitations that may impact its reliability in finding roots. Its effectiveness is heavily based on starting close to a root, and it does not always guarantee convergence when starting from any arbitrary point. There is also a risk of divergence or of converging to a root that was not intended; see the column *x*_0_ = 6 in Table 1. Each run of the Newton-Raphson method can only isolate one root, provides a floating-point approximation without specific error margins, and struggles with singularities where the derivative *f* ^′^(*x*) equals zero. However, interval-based methods such as the Newton method and Krawczyk methods address these problems by operating with intervals, which helps to ensure global convergence. This approach allows for finding all roots within a given search area and manages singularities through the use of interval arithmetic, see Table 1.

**Table 1.** Steady-state calculation using Newton-Raphson and interval methods. *x*_0_: initial guess; *X*_0_: initial enclosure; 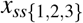: steady states.

|  | Newton-Raphson |  |  |  |  |  |  |  | Interval Newton | Interval Krawczyk |
| --- | --- | --- | --- | --- | --- | --- | --- | --- | --- | --- |
| $x_0$ or $X_0$ | $x_0 = 0$ | $x_0 = 0.1$ | $x_0 = 1$ | $x_0 = 5$ | $x_0 = 6$ | $x_0 = 10$ | $x_0 = 50$ | $x_0 = 99$ | $X_0 = [0, 500]$ | $X_0 = [0, 500]$ |
| $x_{ss1}$ | 0 | 0 | 0 | 0 | Not found | Not Found | Not found | Not found | $[0, 0.0008]$ | $[0, 2.62 \times 10^{-70}]$ |
| $x_{ss2}$ | Not found | Not found | Not found | Not found | Not found | $\approx 7.81$ | Not found | Not found | $[7.81293, 7.81295]$ | $[7.728, 7.899]$ |
| $x_{ss3}$ | Not found | Not found | Not found | Not found | $\approx 100$ | Not found | $\approx 100$ | $\approx 100$ | $[99.9999, 100]$ | $[99.9999, 100]$ |

The same parameter intervals and initial enclosure described in the previous example were used for all interval methods used to determine interval bounds on steady-state solutions. The guaranteed bounds calculated for multiple steady states, derived using the underlying methods, are shown in Fig. 2. It should be noted that interval methods can effectively determine all the roots, including an unstable steady state lying between two stable steady states, a task that is often very challenging for conventional numerical approaches.

**Fig. 2.**
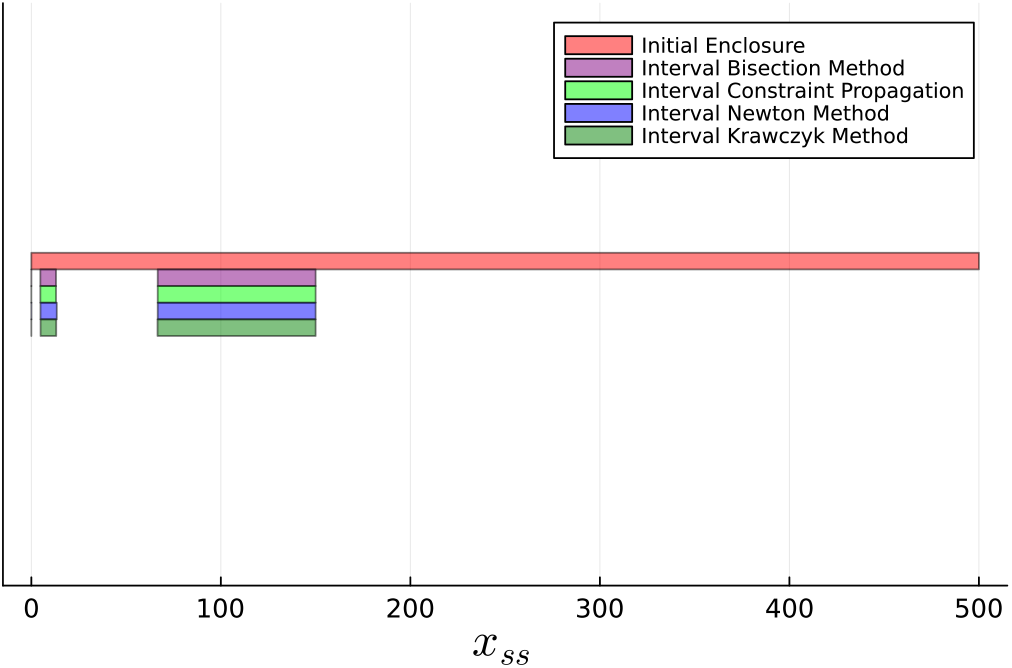
Positive Feedback Loop: Rigorous bounds on three steady states, two stable and one unstable in between, for given parameter intervals using interval methods. Parameter ranges: *α* = [200, 300], *γ* = [2, 3], *K* = [20*/*3, 15]. Initial enclosure of steady states: *X*_0_ = [0, 500].

### 3.3 Dual Positive Feedback Loop

This model represents a dual feedback mechanism (a multiple positive feedback loop) in the galactose regulatory network as given in (Venturelli et al., 2012). This model consists of three coupled equations:

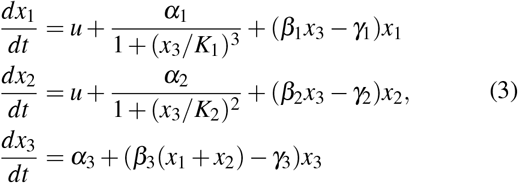

Analysis of this multistable model demonstrates how the interval methods handle nonlinear systems with strong interdependencies. We take *u* = 1, and the parameter values are specified as follows: *α*_1_ = *α*_2_ = *α*_3_ = [95, 105], *γ*_1_ = *γ*_2_ = *γ*_3_ = [0.95, 1.95], *K*_1_ = *K*_2_ = [9.5, 10.5], and *β*_1_ = *β*_2_ = *β*_3_ = and Fig. 4, respectively. [−0.525, −0.475]. The initial enclosure of the solution is given by *X*_0_ = [0, 200]^3^. The graphical representation of the computed bounds, as determined using the interval constraint propagation method and the interval Newton method, is presented in Fig. 3

**Fig. 3.**
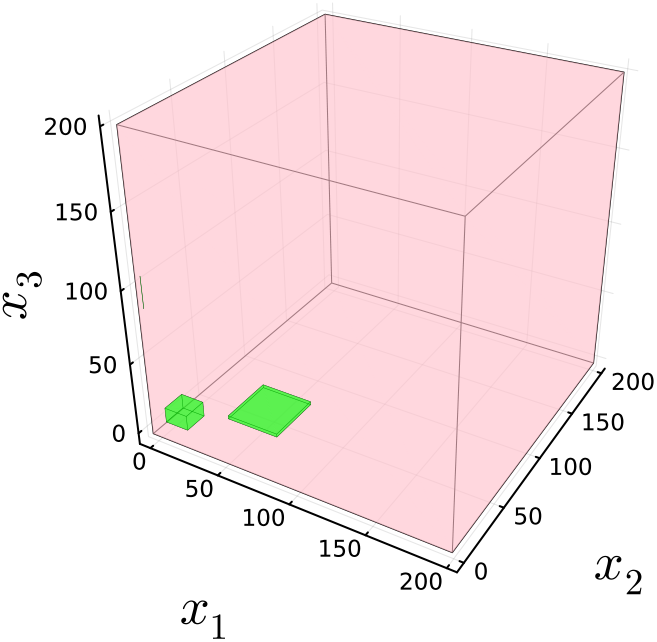
The figure illustrates the use of the interval constraint propagation method applied to a three-dimensional dual positive feedback loop model. Within the specified parameter intervals, the method guarantees enclosure on multiple steady states, as shown by the green boxes. The initial enclosure on the solution is depicted by the pink box. With *u* = 1, the parameter values are: *α*_1_ = *α*_2_ = *α*_3_ = [95, 105], *γ*_1_ = *γ*_2_ = *γ*_3_ = [0.95, 1.95], *K*_1_ = *K*_2_ = [9.5, 10.5], *β*_1_ = *β*_2_ = *β*_3_ = [−0.525, −0.475]. The initial solution enclosure is *X*_0_ = [0, 200]^3^.

**Fig. 4.**
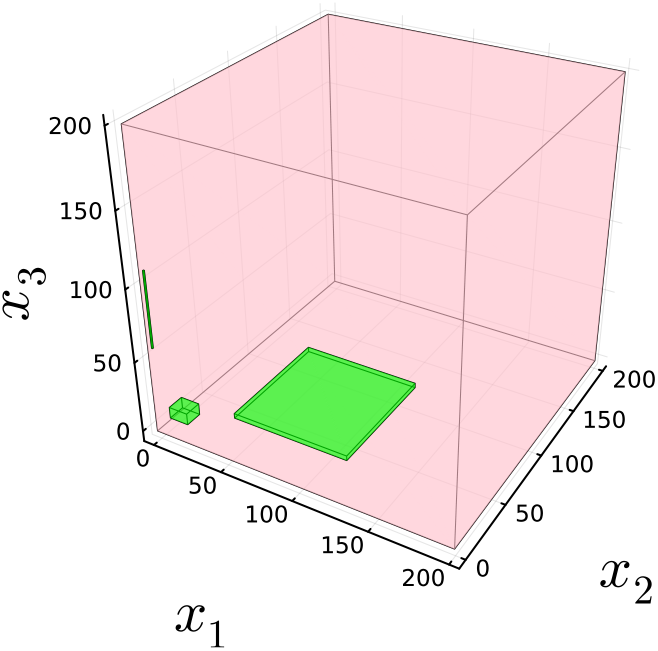
The same analysis as shown in Fig. 3 but using the interval Newton method. The bounds are comparatively conservative than the interval constraint propagation due to ill-conditioned Jacobian matrix.

### 3.4 Incoherent Feedforward Loop

This two-dimensional incoherent feedforward loop model is one of the key network motifs found in complex gene regulatory networks. It is also capable of showcasing a diverse range of dynamic behaviours, such as adaptation, pulsed response, foldchange detection, etc. The mathematical model of an incoherent feedforward loop based on protein interactions (Del Vecchio and Murray, 2015) is shown as

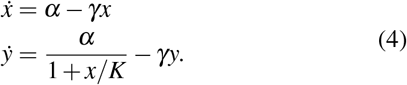

The parameters employed are consistent with those outlined in Example 3.1 and Example 3.2. The initial enclosure is specified as *X*_0_ = [0, 500]^2^. The computation of rigorous bounds on steady states utilising the Krawczyk method is illustrated in Figure 5.

**Fig. 5.**
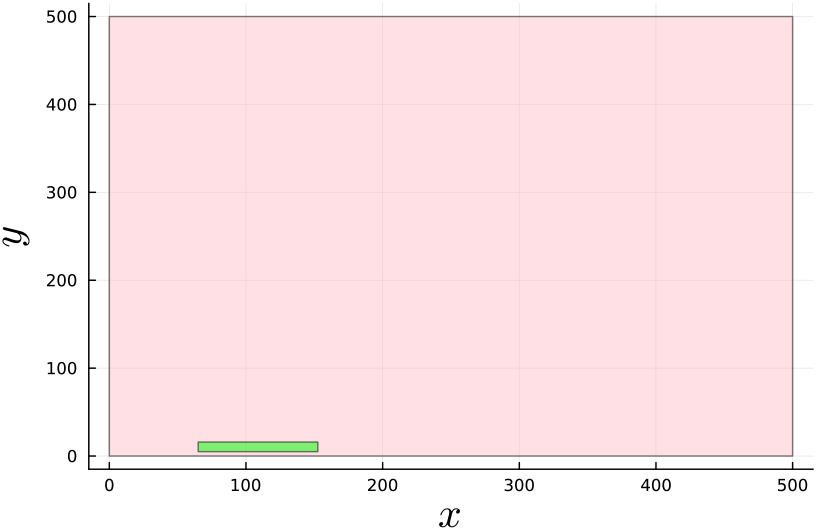
Steady-state bounds (green) for given parameter intervals using the interval Krawczyk method. The initial enclosure is represented by a pink colour box. Parameter ranges: *α* = [200, 300], *γ* = [2, 3], *K* = [20*/*3, 15]. Initial enclosure of steady states: *X*_0_ = [0, 500]^2^.

### 3.5 Synthetic Incoherent Feedforward Loop

This last synthetic circuit example is a three-dimensional systems with nonlinear dynamics, based on design given in (Kim et al., 2014),

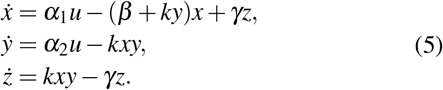

The parameter intervals are specified as follows: *α*_1_ = *α*_2_ = [0.0128, 0.0142], *β* = [0.00171, 0.0019], *γ* = [0.001045, 0.001155], and *k* = [14249.9, 15750.1]. The initial enclosure of the solution is given by *X*_0_ = [0, 5] × [0, 5 × 10^−7^] × [0, 5]. The rigorous enclosure of steady states using the Krawczyk interval method is shown in Figure 6.

**Fig. 6.**
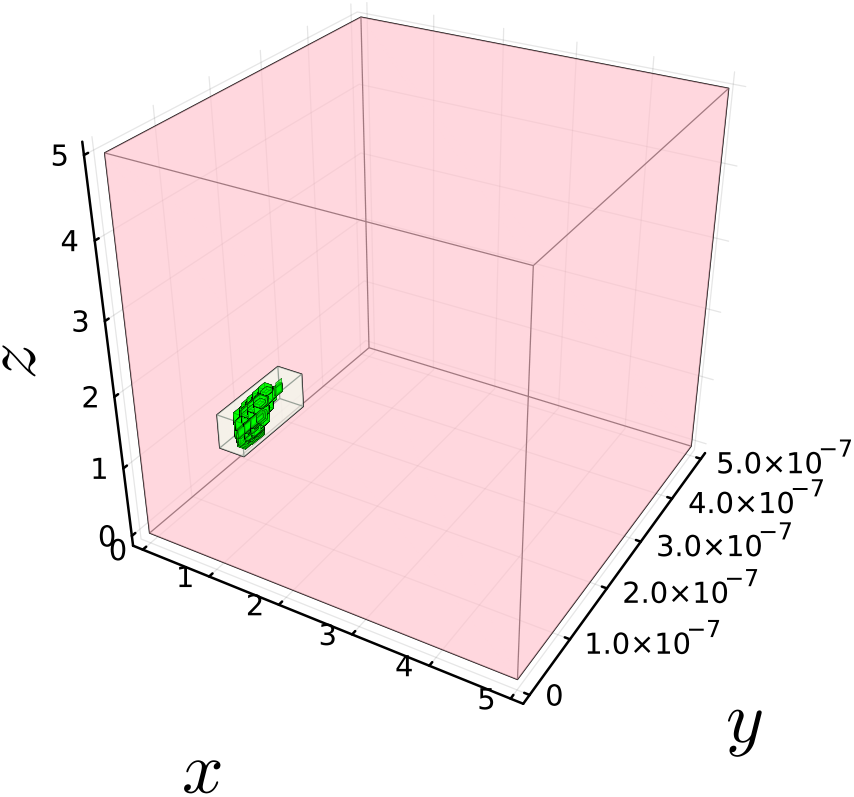
The figure illustrates the bounded steady-state solutions of a three-dimensional nonlinear synthetic incoherent feedforward loop. These bounds are depicted in green, corresponding to given parameter intervals calculated by the interval Krawczyk method. The presence of green subboxes results from a subdivision of the search space to mitigate overestimation caused by interval dependency. Initially, the enclosure within which these computations take place is indicated by a pink-coloured box. The parameter intervals are: *α*_1_ = *α*_2_ = [0.0128, 0.0142], *β* = [0.00171, 0.0019], *γ* = [0.001045, 0.001155], *k* = [14249.9, 15750.1]. The initial solution enclosure is *X*_0_ = [0, 5] × [0, 5 × 10^−7^] × [0, 5].

## 4. COMPARISON

The examples analysed underscore the critical role of interval methods in addressing parametric uncertainties within bounded steady-state computations of nonlinear biomolecular systems. Table 2 presents the computed guaranteed bounds for all the discussed examples utilising all the interval methods discussed in the previous section.

**Table 2.**
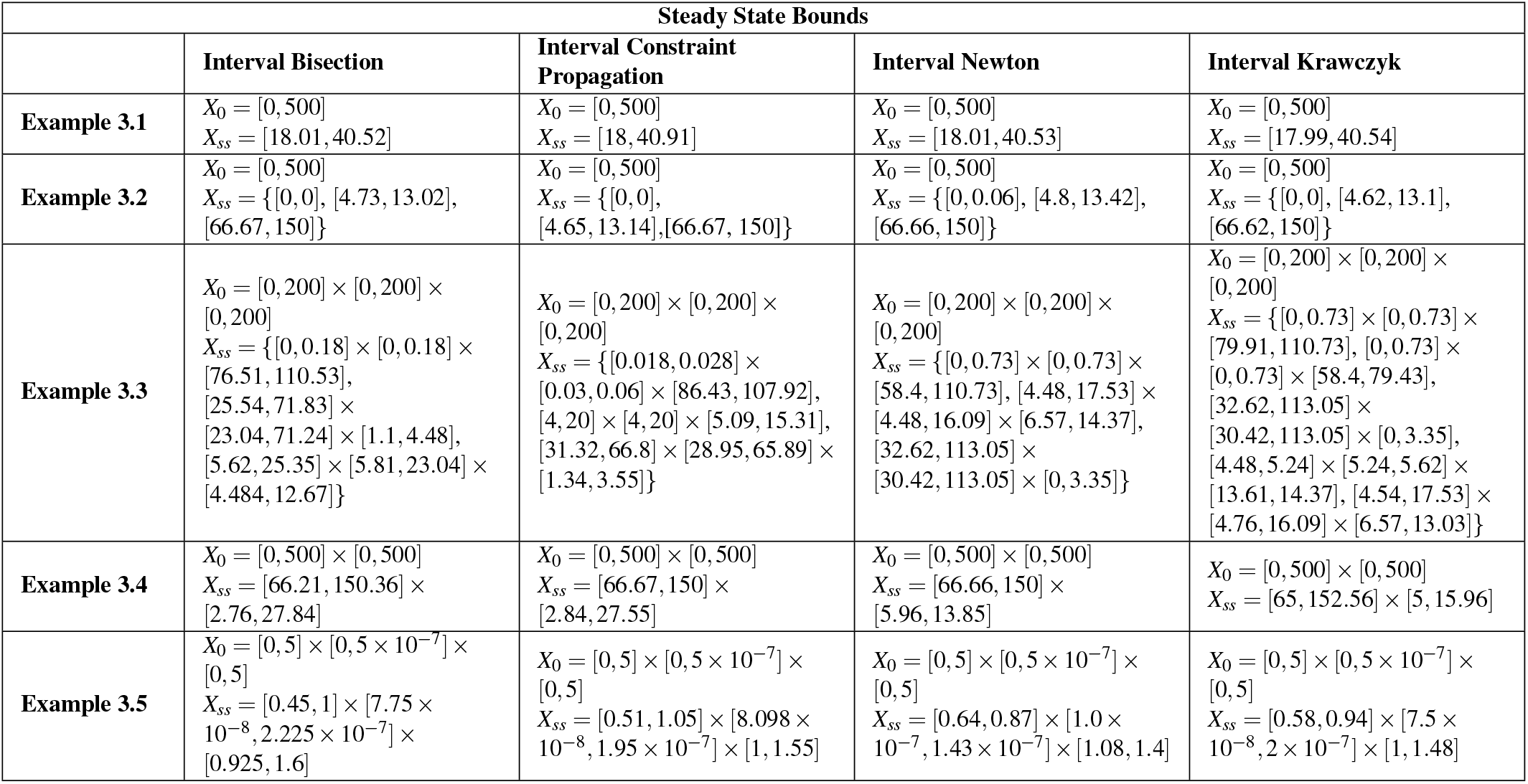
Comparing calculated steady-state bounds using interval methods.

| Steady State Bounds |  |  |  |  |
| --- | --- | --- | --- | --- |
|  | Interval Bisection | Interval Constraint Propagation | Interval Newton | Interval Krawczyk |
| **Example 3.1** | $X_0 = [0, 500]$<br>$X_{ss} = [18.01, 40.52]$ | $X_0 = [0, 500]$<br>$X_{ss} = [18, 40.91]$ | $X_0 = [0, 500]$<br>$X_{ss} = [18.01, 40.53]$ | $X_0 = [0, 500]$<br>$X_{ss} = [17.99, 40.54]$ |
| **Example 3.2** | $X_0 = [0, 500]$<br>$X_{ss} = \{[0, 0], [4.73, 13.02], [66.67, 150]\}$ | $X_0 = [0, 500]$<br>$X_{ss} = \{[0, 0], [4.65, 13.14], [66.67, 150]\}$ | $X_0 = [0, 500]$<br>$X_{ss} = \{[0, 0.06], [4.8, 13.42], [66.66, 150]\}$ | $X_0 = [0, 500]$<br>$X_{ss} = \{[0, 0], [4.62, 13.1], [66.62, 150]\}$ |
| **Example 3.3** | $X_0 = [0, 200] \times [0, 200] \times [0, 200]$<br>$X_{ss} = \{[0, 0.18] \times [0, 0.18] \times [76.51, 110.53], [25.54, 71.83] \times [23.04, 71.24] \times [1.1, 4.48], [5.62, 25.35] \times [5.81, 23.04] \times [4.484, 12.67]\}$ | $X_0 = [0, 200] \times [0, 200] \times [0, 200]$<br>$X_{ss} = \{[0.018, 0.028] \times [0.03, 0.06] \times [86.43, 107.92], [4, 20] \times [4, 20] \times [5.09, 15.31], [31.32, 66.8] \times [28.95, 65.89] \times [1.34, 3.55]\}$ | $X_0 = [0, 200] \times [0, 200] \times [0, 200]$<br>$X_{ss} = \{[0, 0.73] \times [0, 0.73] \times [58.4, 110.73], [4.48, 17.53] \times [4.48, 16.09] \times [6.57, 14.37], [32.62, 113.05] \times [30.42, 113.05] \times [0, 3.35]\}$ | $X_0 = [0, 200] \times [0, 200] \times [0, 200]$<br>$X_{ss} = \{[0, 0.73] \times [0, 0.73] \times [79.91, 110.73], [0, 0.73] \times [0, 0.73] \times [58.4, 79.43], [32.62, 113.05] \times [30.42, 113.05] \times [0, 3.35], [4.48, 5.24] \times [5.24, 5.62] \times [13.61, 14.37], [4.54, 17.53] \times [4.76, 16.09] \times [6.57, 13.03]\}$ |
| **Example 3.4** | $X_0 = [0, 500] \times [0, 500]$<br>$X_{ss} = [66.21, 150.36] \times [2.76, 27.84]$ | $X_0 = [0, 500] \times [0, 500]$<br>$X_{ss} = [66.67, 150] \times [2.84, 27.55]$ | $X_0 = [0, 500] \times [0, 500]$<br>$X_{ss} = [66.66, 150] \times [5.96, 13.85]$ | $X_0 = [0, 500] \times [0, 500]$<br>$X_{ss} = [65, 152.56] \times [5, 15.96]$ |
| **Example 3.5** | $X_0 = [0, 5] \times [0, 5 \times 10^{-7}] \times [0, 5]$<br>$X_{ss} = [0.45, 1] \times [7.75 \times 10^{-8}, 2.225 \times 10^{-7}] \times [0.925, 1.6]$ | $X_0 = [0, 5] \times [0, 5 \times 10^{-7}] \times [0, 5]$<br>$X_{ss} = [0.51, 1.05] \times [8.098 \times 10^{-8}, 1.95 \times 10^{-7}] \times [1, 1.55]$ | $X_0 = [0, 5] \times [0, 5 \times 10^{-7}] \times [0, 5]$<br>$X_{ss} = [0.64, 0.87] \times [1.0 \times 10^{-7}, 1.43 \times 10^{-7}] \times [1.08, 1.4]$ | $X_0 = [0, 5] \times [0, 5 \times 10^{-7}] \times [0, 5]$<br>$X_{ss} = [0.58, 0.94] \times [7.5 \times 10^{-8}, 2 \times 10^{-7}] \times [1, 1.48]$ |

The interval bisection method effectively eliminates nonsolution regions of the search space but can be computationally intensive due to its recursive nature. In contrast, the interval constraint propagation method achieves this reduction more efficiently by employing the set-inversion approach. However, in scenarios involving multistability, the interval bisection or sub-division + filter method is additionally required. The interval Newton method offers quadratic convergence, ensuring rapid convergence to the solution, though it may be less effective for ill-conditioned systems where the inverse of the Jacobian is undefined for a given interval. The interval Krawczyk method effectively resolves this issue by offering a robust solution approach for ill-conditioned systems, as it eliminates the necessity of computing the inverse of the interval matrix. Based on a comprehensive analysis of biomolecular systems using interval methods, our recommendations on the appropriate method for various types of problem are presented in Fig. 7.

**Fig. 7.**
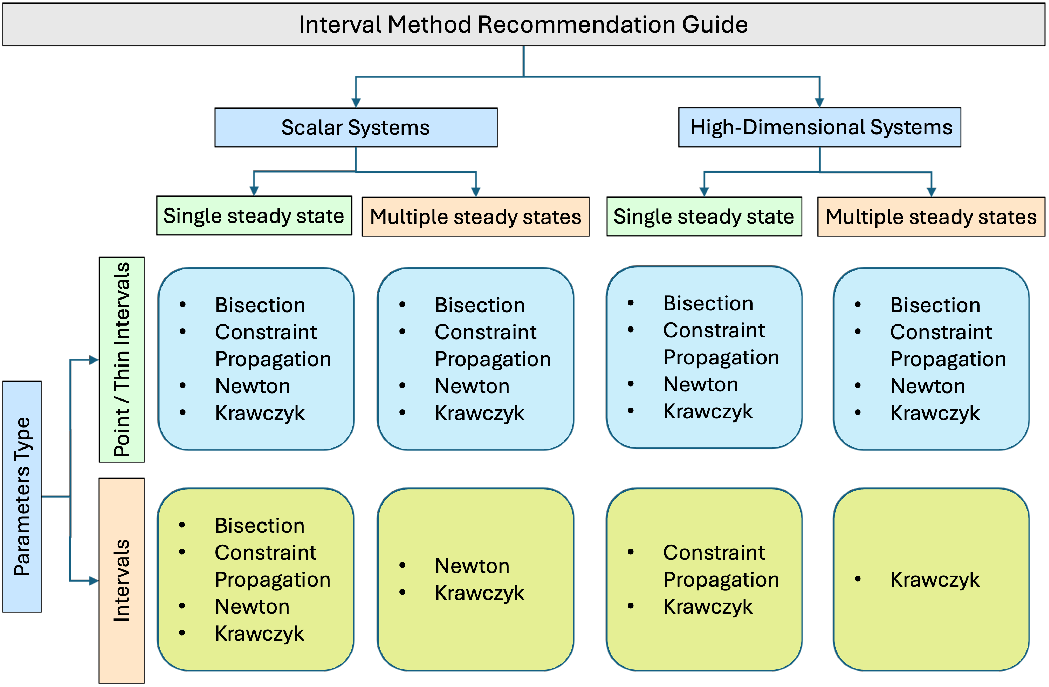
This figure illustrates our recommendation of which method we should choose in which type of problem. A thin interval involves expressing a point value within an interval to account for the rounding errors associated with floating-point numbers, for example, 1 ∈ [0.999, 1.001].

## 5. CONCLUSION

The paper addresses the problem of globally determining guaranteed solutions for complex, multidimensional biomolecular systems that exhibit multiple stable states and contain uncertain parameters. It presents algorithms for the interval Newton and Krawczyk methods, which are used to find roots in these complicated systems. The study compares four different interval methods: interval bisection, interval constraint propagation, interval Newton, and interval Krawczyk. It assesses how rigorous and effective these methods are in bounding the steadystate solutions of uncertain, multistable, and multidimensional nonlinear systems.

The interval Newton method is noted for quadratic convergence in non-singular regimes, while the Krawczyk method excels in ill-conditioned multidimensional systems without needing the inverse of an interval matrix.

Based on this analysis, recommendations are provided for the applicability of the method to specific problems. Interval Constraint Propagation is highlighted for its efficiency in multiple steady states by using set-inversion with bisection strategies in multistable scenarios. Numerical examples, including biologically relevant models, demonstrate the framework’s capability in handling multistability and interval dependency. The study extends the methodologies to include parameter uncertainty, showing robust enclosures of steady states amid biological variability, effectively identifying stable and unstable states. Suggesting parameter subdivision and hybrid strategies, the paper argues for improved speed and reliability by reducing interval dependency.

This study offers four pivotal advancements. First, it involves the refinement of current numerical techniques to effectively analyse nonlinear models of biomolecular circuits. Second, the study broadens existing methodologies to address parameter uncertainty, thereby providing robust encapsulations of steady states amidst significant biological variability. Third, it merges validated numerical methods with systems biology to establish a dependable and comprehensive framework for rigorous steady-state analysis. This integration improves the predictability and reliability of therapeutic interventions and strategies even under high uncertainty, directly contributing to patient safety and minimising risks in drug discovery and therapeutic design. Lastly, the research supports the scalable analysis of complex systems and the robust design of synthetic circuits, presenting valuable tools for advances in synthetic biology, gene editing, and molecular engineering. For example, bistable transcriptional switches depend on precise sequestration dynamics to ensure reliable switching (Kim et al., 2006). On the other hand, bistable circuits based on CRISPR interference (CRISPRi) and dCas9 achieve stable expression states through mutual repression (Santos-Moreno et al., 2020).

Future research should focus on improving computational efficiency, especially in high dimensions, and developing strategies to reduce interval overestimation with hybrid approaches.

## Notes

### Competing Interest Statement

The authors have declared no competing interest.

### Summary of Updates

This version of the manuscript has been revised to (i) add S. Janardhanan to the author list and (ii) include an IFAC acceptance and licensing preprint notice (Creative Commons CC-BY-NC-ND) in the title/footnote section.

## REFERENCES

(2025). Code for examples. https://github.com/RudraPrakashIITDelhi/ BioSystems-Rigorous-Analysis. Accessed: 25 November 2025.

Alon, U. (2006). An Introduction to Systems Biology: Design Principles of Biological Circuits. Chapman and Hall/CRC, New York.

Angeli, D., Ferrell, J.E., and Sontag, E.D. (2004). Detection of multistability, bifurcations, and hysteresis in a large class of biological positive-feedback systems. Proceedings of the National Academy of Sciences of the United States of America, 101(7), 1822–1827.

Avcu, N. and Güzeliş, C. (2019). Bifurcation analysis of bistable and oscillatory dynamics in biological networks using the root-locus method. IET Systems Biology, 13(6), 333–345. Publisher: The Institution of Engineering and Technology.

Bagci, E.Z., Vodovotz, Y., Billiar, T.R., Ermentrout, G.B., and Bahar, I. (2006). Bistability in Apoptosis: Roles of Bax, Bcl-2, and Mitochondrial Permeability Transition Pores. Biophysical Journal, 90(5), 1546–1559.

Chorasiya, G., Prakash, R., and Sen, S. (2023). Quantitative Steady-State Bounds in Biomolecular Circuits Due to Bounded Multi-Parametric Perturbations. IEEE Control Systems Letters, 7, 2827–2832.

Del Vecchio, D. and Murray, R.M. (2015). Biomolecular feedback systems. Princeton University Press Princeton, NJ.

Dey, A. and Barik, D. (2021). Potential Landscapes, Bifurcations, and Robustness of Tristable Networks. ACS Synthetic Biology, 10(2), 391–401. Publisher: American Chemical Society.

Ferrell, J.E. (2002). Self-perpetuating states in signal transduction: positive feedback, double-negative feedback and bistability. Current Opinion in Cell Biology, 14(2), 140–148.

Filippi, S., Barnes, C.P., Kirk, P.D.W., Kudo, T., Kunida, K., McMahon, S.S., Tsuchiya, T., Wada, T., Kuroda, S., and Stumpf, M.P.H. (2016). Robustness of MEK-ERK Dynamics and Origins of Cell-to-Cell Variability in MAPK Signaling. Cell Reports, 15(11), 2524–2535.

Gardner, T.S., Cantor, C.R., and Collins, J.J. (2000). Construction of a genetic toggle switch in Escherichia coli. Nature, 403(6767), 339–342. Publisher: Nature Publishing Group.

Hasenauer, J., Rumschinski, P., Waldherr, S., Borchers, S., Allgöwer, F., and Findeisen, R. (2009). Guaranteed Steady-State Bounds for Uncertain Chemical Processes. IFAC Proceedings Volumes, 42(11), 643–648.

Jaulin, L. (2000). Interval constraint propagation with application to bounded-error estimation. Automatica, 36(10), 1547–1552. Publisher: Elsevier.

Jaulin, L., Kieffer, M., Didrit, O., and Walter, E. (2001). Applied Interval Analysis. Springer London, London.

Kim, J., Khetarpal, I., Sen, S., and Murray, R.M. (2014). Synthetic circuit for exact adaptation and fold-change detection. Nucleic Acids Research, 42(9), 6078–6089.

Kim, J., White, K.S., and Winfree, E. (2006). Construction of an in vitro bistable circuit from synthetic transcriptional switches. Molecular Systems Biology, 2, 68.

Kuntz, J., Thomas, P., Stan, G.B., and Barahona, M. (2019). Bounding the stationary distributions of the chemical master equation via mathematical programming. The Journal of Chemical Physics, 151(3), 34109.

Ma, W., Trusina, A., El-Samad, H., Lim, W.A., and Tang, C. (2009). Defining Network Topologies that Can Achieve Biochemical Adaptation. Cell, 138(4), 760–773.

Moore, R.E. (1977). A test for existence of solutions to nonlinear systems. SIAM Journal on Numerical Analysis, 14(4), 611–615. Publisher: SIAM.

Moore, R.E., Kearfott, R.B., and Cloud, M.J. (2009). Introduction to interval analysis. SIAM.

Neumaier, A. (1990). Interval Methods for Systems of Equations. Cambridge University Press.

Otero-Muras, I., Banga, J.R., and Alonso, A.A. (2012). Characterizing Multistationarity Regimes in Biochemical Reaction Networks.

Prakash, R., Janardhanan, S., and Sen, S. (2024). Design of parameter intervals to meet steady-state specifications in biomolecular circuits using interval analysis. bioRxiv, 2024–10. doi:10.1101/2024.10.24.620126.

Reyes, B.C., Otero-Muras, I., and Petyuk, V.A. (2022). A numerical approach for detecting switch-like bistability in mass action chemical reaction networks with conservation laws. BMC Bioinformatics, 23(1), 1.

Rump, S.M. (2010). Verification methods: rigorous results using floating-point arithmetic. In Proceedings of the 2010 International Symposium on Symbolic and Algebraic Computation, 3–4. ACM, Munich Germany.

Santos-Moreno, J., Tasiudi, E., Stelling, J., and Schaerli, Y. (2020). Multistable and dynamic CRISPRi-based synthetic circuits. Nature Communications, 11(1), 2746. Publisher: Nature Publishing Group.

Sreedharan, V., Bhalla, U.S., and Ramakrishnan, N. (2023). Using sensitivity analyses to understand bistable system behavior. BMC Bioinformatics, 24(1), 136.

Tucker, W. (2011). Validated numerics. In Validated Numerics. Princeton University Press.

Venturelli, O.S., El-Samad, H., and Murray, R.M. (2012). Synergistic dual positive feedback loops established by molecular sequestration generate robust bimodal response. Proceedings of the National Academy of Sciences, 109(48).

